# Commitments by the Biopharmaceutical Industry to Clinical Trial Transparency: The Evolving Environment

**DOI:** 10.1101/349902

**Authors:** Slavka Baronikova, Jim Purvis, Eric Southam, Julie Beeso, Christopher C Winchester, Antonia Panayi

## Abstract

**Background:** Sponsors of clinical trials have ethical obligations to register protocols, to report study results and to comply with applicable legal requirements.

**Objective:** To evaluate public commitments to trial disclosure and rates of disclosure by members and non-members of the European Federation of Pharmaceutical Industries and Associations (EFPIA) and/or the Pharmaceutical Research and Manufacturers of America (PhRMA).

**Study selection:** Websites of the top 50 biopharmaceutical companies by 2015 sales were searched for statements relating to trial data disclosure. Disclosure of trial results completed by biopharmaceutical industry and non-industry sponsors of at least 30 trials (2006–2015) was assessed using TrialsTracker.

**Findings:** Among the top 50 companies, 30 were EFPIA/PhRMA members and 20 were non-members, of which 26 and none, respectively, had a statement on their website committing to the disclosure of trials data. Of 29 377 trials in TrialsTracker, 9511 were industry-sponsored (69 companies) and 19 866 were non-industry-sponsored (254 institutions). The overall mean disclosure rate was 55%, with higher rates for industry (74%) than for non-industry sponsors (46%). Of the 30 companies within the top 50 with data in TrialsTracker, the mean disclosure rate was 76% (77% for EFPIA/PhRMA members [n = 25] versus 67% for non-members [n = 5]).

**Conclusions:** Most of the top 50 biopharmaceutical companies have publicly committed to the disclosure of trial data. Industry sponsors have responded to the ethical and legal demands of trial disclosure to a greater extent than non-industry sponsors, and now disclose three quarters of their trials.

## INTRODUCTION

A perceived lack of transparency, including under-reporting of results, undermines the confidence of researchers, healthcare professionals and patients in conclusions drawn from clinical trials.[1] All clinical trial sponsors, be they biopharmaceutical companies or non-industry bodies, such as government agencies, universities and research charities, have ethical obligations to register trials before they start and to report their results in a timely fashion after they finish.[2,3] In the USA, EU and elsewhere, it is required that certain types of clinical trial are registered and their results posted on dedicated registries (e.g. EudraCT, the EU electronic Register of Post-Authorisation Studies and ClinicalTrials.gov) (Supplementary material, table S1).[4-11] Other bodies, such as the World Health Organization (WHO) and the International Committee of Medical Journal Editors (ICMJE), have issued transparency standards and recommendations,[2,12-14] and some biopharmaceutical companies have websites dedicated to their own trial results.[15,16] This makes the clinical trial data transparency environment highly complex and diverse.

Within the biopharmaceutical industry, which is responsible for approximately half of all clinical trials,[17,18] two large associations, the European Federation of Pharmaceutical Industries and Associations (EFPIA) and the Pharmaceutical Research and Manufacturers of America (PhRMA), have developed joint “Principles for responsible clinical trial data sharing”.[19] These joint principles, which became effective on 1 January 2014, make the following five commitments:

1. to enhance data sharing with researchers
2. to enhance public access to clinical study information
3. to share results with patients who participate in clinical trials
4. to certify procedures for disclosing clinical trial information
5. to reaffirm commitments to publish clinical trial results.

In the present study, we aimed to evaluate the extent to which EFPIA/PhRMA members and non-members among the leading biopharmaceutical companies have committed to the responsible disclosure of clinical trial results. We also evaluated the reporting of results from clinical trials sponsored by biopharmaceutical companies compared with those from other sponsors.

## METHODS

### Commitment to disclosure of clinical trial data by EFPIA/PhRMA member companies

The global public websites of each EFPIA and/or PhRMA (‘EFPIA/PhRMA’) member and non-member company in the top 50 companies by 2015 worldwide prescription sales (‘top 50 companies’)[20] were searched by one researcher (JP) for direct links to pages containing: (i) a general statement of commitment to disclosing clinical trial data; (ii) a general statement of commitment to disclosing clinical trial data according to EFPIA/PhRMA joint principles; and (iii) specific statements detailing commitments to upholding one or more of the five individual EFPIA/PhRMA joint principles for responsible disclosure of clinical trial data. If no direct links to such pages were found, the free-text search function of each website was used to search for statements relating to clinical trial data disclosure and implementation of the EFPIA/PhRMA disclosure principles using one or more the key words “EFPIA”, “PhRMA”, “data sharing”, “clinical trials” and “transparency”. EFPIA/PhRMA membership was determined from the websites of these two organisations (www.efpia.eu/about-us/membership and http://www.phrma.org/about/members).

Ease of access to relevant information was assessed; good access was rated as requiring either no more than four clicks from the homepage of the company website, [21] or a clear, direct link, poor access was rated as either needing more than four clicks or requiring navigation to satellite websites (e.g. blogs).

### Clinical trial results reporting

TrialsTracker is an independent, semi-automated, web-based tool that has been developed in an effort to incentivise sponsors of clinical trials to improve disclosure rates by highlighting the disclosure performance of individual sponsors (trials without results disclosed as a proportion of trials registered).[22] For clinical trial sponsors to be included in TrialsTracker, they must have more than 30 phase II–IV clinical trials registered on ClinicalTrials.gov that were recorded as completed after 1 January 2006 and at least 24 months before the most recent TrialsTracker update. Because the most recent update to the database was in April 2017, the most recent studies to be included in this analysis were completed in April 2015.

Data detailing the number of trials registered on ClinicalTrials.gov by clinical trial sponsor, and the corresponding number of trials without results reported for each year from 2006 to 2015, were downloaded as a comma-separated values (CSV) file from the TrialsTracker website (https://trialstracker.ebmdatalab.net). TrialsTracker identifies sponsors as industry (“biopharmaceutical companies” which we sub-categorised as pharmaceutical/biotechnology, generics/biosimilars, medical devices, plasma products and nutraceuticals, using information on the company websites that was found during the research for our study) or non-industry (classified as National Institutes of Health, US Federal or other) institutions. For each industry and non-industry sponsor and for each category, the number of disclosed trials and the percentage of eligible studies with disclosed results were calculated in Microsoft Excel 2016.

An analysis of disclosure rates was performed on subsets of the industry sponsors within TrialsTracker based on sales revenue (the top 50 companies)[20] and membership of EFPIA/PhRMA. An arbitrary disclosure rate threshold of 80% was applied to sponsor subgroups.

### Exploratory analysis of results posted on websites other than ClinicalTrials.gov

In an exploratory analysis, clinical trial results from locations other than ClinicalTrials.gov or from linked publications in PubMed were sought for three studies that were blindly selected from four of the top 50 companies. ClinicalTrials.gov was searched by National Clinical Trial (NCT) identifier in order to establish the presence or absence of posted results and/or links to publications on PubMed. We made a separate search of PubMed, Google Scholar and Google using the NCT identifier, and a search of EudraCT and the relevant company’s website based on NCT identifier and study title.

### Exploratory analysis of commitments to disclosure of clinical trial data by non-industry sponsors

In an exploratory analysis, we searched for statements relating to the disclosure of clinical trial data on the websites of 10 non-industry sponsors of clinical trials with results completed in the period 2006–2015.

### Data analysis

Disclosure rates for all industry sponsors, the top 50 companies and EFPIA/PhRMA members in the top 50 companies were compared with those for non-industry sponsors.

### Patient involvement

Patients were not directly involved in conducting this study, but patients’ perspectives were sought during the development of the manuscript.

## RESULTS

### Commitment to disclosure of clinical trial data

#### EFPIA/PhRMA membership

Of the top 50 companies, six were EFPIA members only, two were PhRMA members only, 22 were both EFPIA and PhRMA members and 20 were neither EFPIA nor PhRMA members. Of the 25 largest companies, 23 were EFPIA/PhRMA members compared with seven of the next largest 25 companies.

All 30 EFPIA/PhRMA members in the top 50 companies were pharmaceutical/biotechnology companies, whereas the 20 non-members were more varied comprised pharmaceutical/biotechnology (n = 8), generics/biosimilars (n = 6), medical devices (n = 1), both generics and medical devices (n = 2), intravenous products and medical devices (n = 1), plasma products (n = 1), and nutraceuticals (n = 1) companies. Of the two EFPIA/PhRMA non-members in the top 25 companies, one was a biotechnology company and the other was a generics company.

#### Access to a general disclosure statement

A general statement committing to the disclosure of clinical trial information was found on 26 of the top 50 company websites (52%), all of which were EFPIA/PhRMA members (table 1). In 19 cases (38% of the top 50 companies; 63% of EFPIA/PhRMA members), the statement was found within four clicks of entering the website; in seven cases, access was rated as poor. An analysis of the proportion of EFPIA/PhRMA members versus non-members in the top 50 companies with statements committing to responsible data transparency is shown in Supplementary material, table S2.

**Table 1.** Number of EFPIA and PhRMA member and non-member companies in the top 50 biopharmaceutical companies as ranked by 2015 worldwide prescription sales and their public commitment to disclosing clinical trial data.

| Top 50 companies: membership of EFPIA and/or PhRMA | General data sharing statement | EFPIA/PhRMA principles | Five EFPIA/PhRMA joint principles of responsible clinical data sharing |  |  |  |  | All five principles |
| --- | --- | --- | --- | --- | --- | --- | --- | --- |
|  |  |  | Commitment to sharing clinical trial data with researchers | Public availability of CSR synopsis as a minimum | Availability of results for trial participants | Public certification of adoption of EFPIA/PhRMA commitments | Commitment to publish clinical trial data (phase III minimum) |  |
| **Member (n = 30)** | 26<br>(86.7%) | 20<br>(66.7%) | 25<br>(83.3%) | 22<br>(73.3%) | 18<br>(60.0%) | 21<br>(70.0%) | 24<br>(80.0%) | 16<br>(53.3%) |
| **Non-member (n = 20)** | 0<br>(0.0%) | 0<br>(0.0%) | 0<br>(0.0%) | 1<br>(5.0%) | 0<br>(0.0%) | 0<br>(0.0%) | 0<br>(0.0%) | 0<br>(0.0%) |
| **Total (n = 50)** | 26<br>(52.0%) | 20<br>(40.0%) | 25<br>(50.0%) | 23<br>(46.0%) | 18<br>(36.0%) | 21<br>(42.0%) | 24<br>(48.0%) | 16<br>(32.0%) |

CSR, clinical study report; EFPIA, European Federation of Pharmaceutical Industries and Associations; PhRMA, Pharmaceutical Research and Manufacturers of America.

#### Specific EFPIA/PhRMA principles

An overview statement referring to the adoption of the joint principles was found on 20/30 websites (67%) of EFPIA/PhRMA members. Reference to all five joint principles was found for 16/30 members (53%). Of non-member companies, only one company made a specific disclosure statement (to enhance public access to clinical study information by making synopses of clinical study reports publicly available) (table 1). The most frequently communicated individual commitments were to share clinical trial data with researchers (83% of EFPIA/PhRMA members; 50% of the top 50 companies) and to publish clinical trial data (80% of members; 48% of the top 50 companies).

### Clinical trial results reporting

Of 29 377 trials listed in TrialsTracker, 9511 (32%) were sponsored by 69 biopharmaceutical companies (a mean of 138 trials per company) and 19 866 (68%) were sponsored by 254 non-industry institutions (a mean of 78 trials per institution) (figure 1). Of all undisclosed trials, 81% were sponsored by non-industry institutions and 19% were sponsored by industry. The mean disclosure rate for all trials was 55%, with higher rates for industry (74%) than for non-industry sponsors (46%) (figure 2; Supplementary material, table S2).

**Fig 1.**
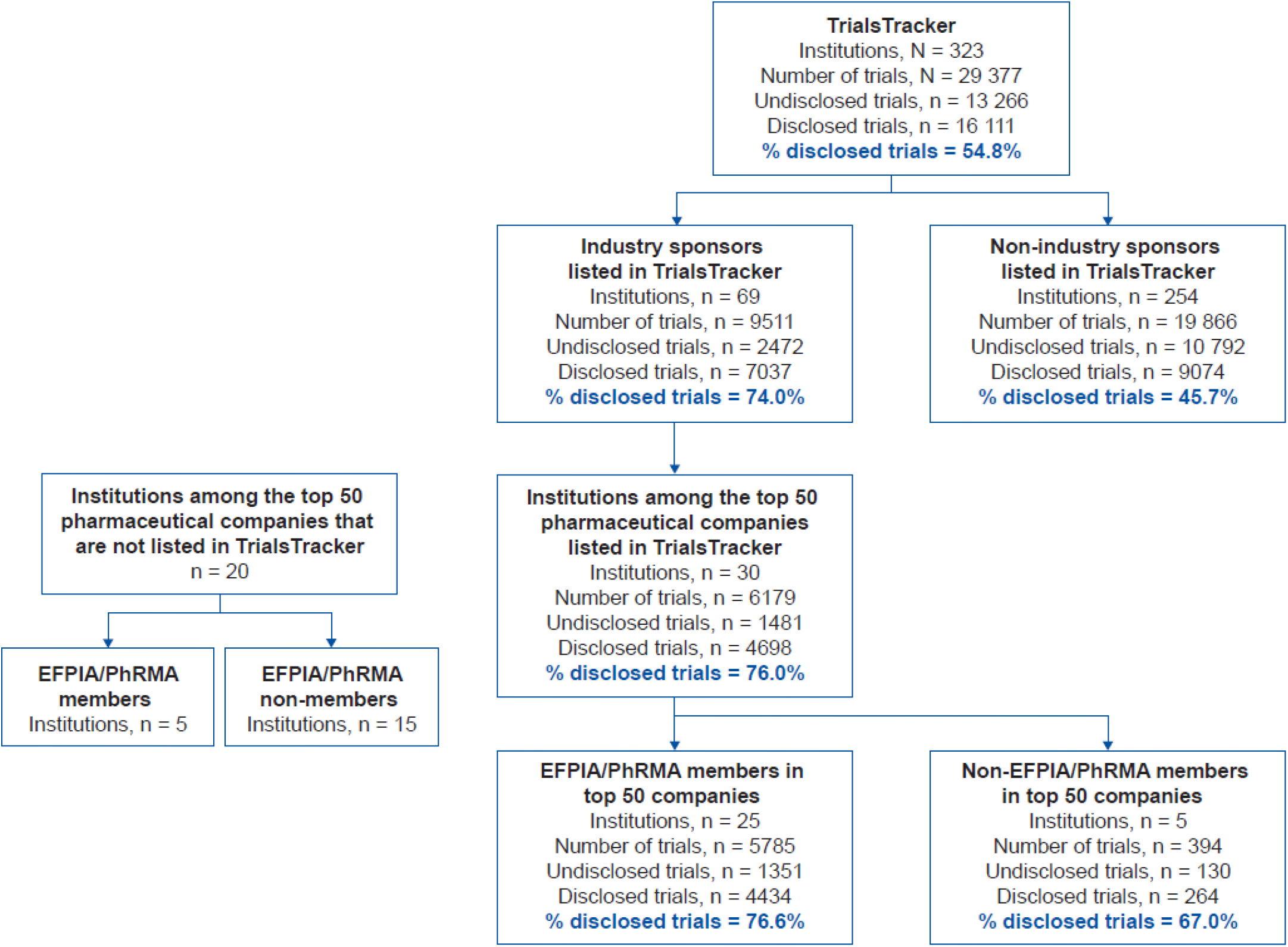
Types and numbers of sponsors represented in TrialsTracker, with their disclosure rates. EFPIA, European Federation of Pharmaceutical Industries and Associations; PhRMA, Pharmaceutical Research and Manufacturers of America.

**Fig 2.**
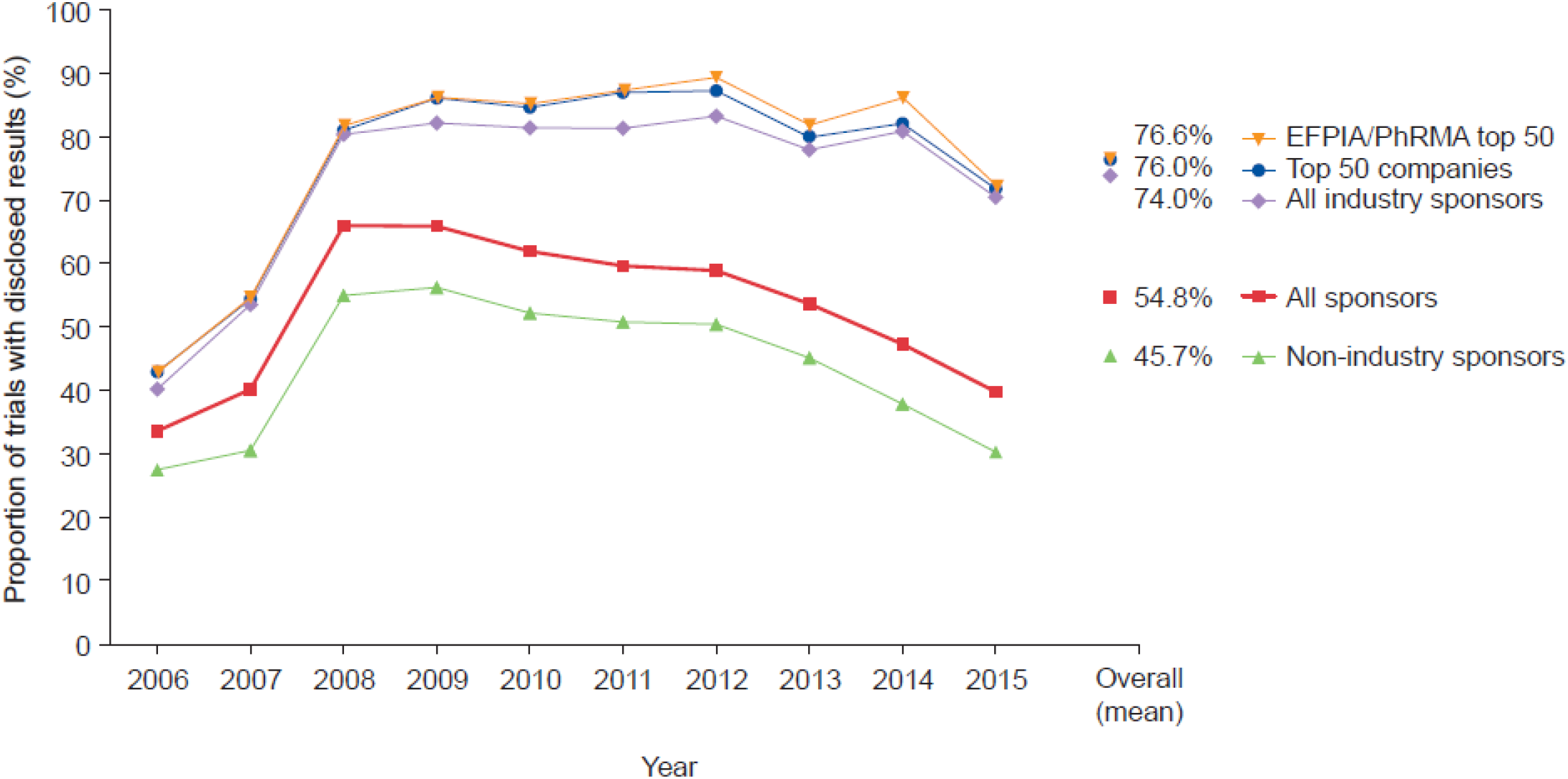
Disclosure of clinical trial results by sponsor type. EFPIA, European Federation of Pharmaceutical Industries and Associations; PhRMA, Pharmaceutical Research and Manufacturers of America.

The overall disclosure rate for all clinical trials substantially increased during 2007 and 2008, before declining thereafter. The maximum mean disclosure rate for all clinical trials was observed in 2008 (66%); for industry- and non-industry-sponsored trials, the maximum mean disclosure rates were in 2012 (83%) and 2009 (56%), respectively (figure 2). Disclosure rates for non-industry sponsors followed a trend similar to those for all sponsors, whereas disclosure rates for industry sponsors were maintained at approximately 80% until 2014 (figure 2).

There was high variability in disclosure rate between sponsor type (figure 3A). The highest disclosure rate achieved by a non-industry sponsor was 84%, whereas two biopharmaceutical industry sponsors achieved 100% disclosure. Of the top 50 companies, a mean of 76% of trials were disclosed by the 30 companies with data reported in TrialsTracker (all of which were pharmaceutical/biotechnology companies). The mean disclosure rate was 77% for EFPIA/PhRMA members (25 companies) and was 67% for non-members (5 companies) (figures 2 and 3B). An arbitrary disclosure rate threshold of 80% was reached by fewer than 1% of non-industry sponsors compared with 39% of industry sponsors. Of the 69 biopharmaceutical industry sponsors with results in TrialsTracker, the 80% threshold was met by 56% of EFPIA/PhRMA members in the top 50 companies, by 20% of EFPIA/PhRMA non-members in the top 50 companies, and by 31% of biopharmaceutical industry sponsors that were not in the top 50 companies (figure 3B).

**Fig 3.**
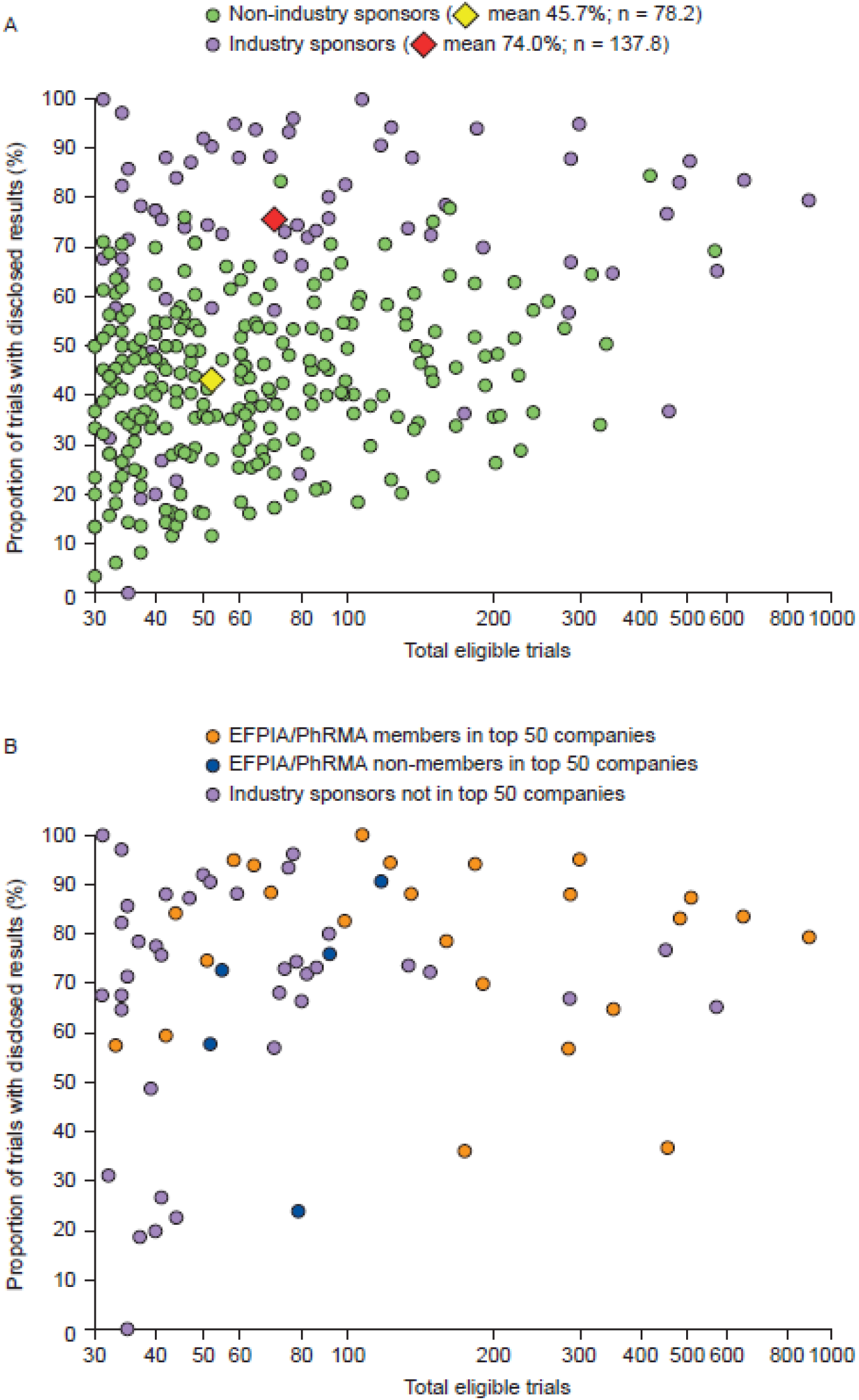
Disclosure rates versus total eligible trials for (A) biopharmaceutical industry and non-industry clinical trial sponsors and (B) industry-only sponsors, highlighting EFPIA/PhRMA members and non-members in the top 50 companies, and industry sponsors not in the top 50 companies. EFPIA, European Federation of Pharmaceutical Industries and Associations; PhRMA, Pharmaceutical Research and Manufacturers of America.

## DISCUSSION

This analysis of the disclosure environment of clinical trial sponsors began with a review of the publicly stated disclosure policies of the top 50 biopharmaceutical companies. Of these, 26 companies (52%; all of which were members of one or both of the two leading international industry bodies [EFPIA and PhRMA]) had disclosure policies available on their websites. Most EFPIA/PhRMA members (87%) communicated that they had a commitment to disclose clinical trial results and two-thirds (67%) specifically referred to the EFPIA/PhRMA joint principles; approximately half (53%) described those principles in detail.

To be useful, information on websites should be easy to find and have a logical flow. The ‘three-click rule’ is no longer regarded as the benchmark for website utility,[21,23] therefore four clicks were used to define information as having good access. Inherent in the principle of publicly committing to data disclosure should be that the statements have good accessibility.

The second phase of our collated disclosure information from a large number of trials and sponsors over a 10-year period using data from TrialsTracker.[22] This showed that the disclosure of clinical trial data remains suboptimal. By the end of April 2017, data was disclosed for approximately half of the phase II–IV trials registered on ClinicalTrials.gov and completed in the period 2006–2015. Over this period, results were disclosed by approximately three-quarters of the biopharmaceutical industry sponsors compared with less than half of the non-industry sponsors. Of undisclosed trials, more than 80% were originally funded by governmental, charitable or academic institutions compared with just under 20% by industry, even though approximately one-third of all trials in our data set were industry funded. Disclosure rates for both types of funder substantially increased between 2007 and 2008, coinciding with mandatory reporting as required by the Food and Drug Administration Amendments Act 801 (FDAAA 801). For industry sponsors, disclosure rates were maintained for the next 6 years before declining slightly in 2015. This decline may reflect delays in the publication process, which usually takes approximately 2 years from study completion.[24] With the implementation of the ‘Final Rule’ in January 2018, there should no longer be delays in the posting of results from applicable clinical trials.[24-26] By contrast, disclosure rates for non-industry studies steadily declined after 2009, possibly reflecting a lack of policing of FDAAA 801.

The proportion of trials identified in the present study as sponsored by the biopharmaceutical industry (approximately one-third of all trials) was similar to that reported previously.[17,26] Although the disclosure rate for industry-sponsored studies was similar to that previously observed using TrialsTracker and the EU Clinical Trials Register (EUCTR),[22,27] this rate is lower than the rates reported for newly approved drugs in the USA and Europe,[28,29] and either lower than or similar to rates of publication that have been reported by single sponsors.[24,25] Similarly, for non-industry studies, the disclosure rate was similar to that previously seen with TrialsTracker,[22] and either lower than or similar to those reported for academic medical centres in the USA and the UK, but higher than for EUCTR.[18,27,30,31]

Our assessment of biopharmaceutical company disclosure policies showed results similar to those from a recent EFPIA/PhRMA survey in which 77% of the 44 EFPIA/PhRMA members confirmed that they state on a publicly available website that they adhere to the joint principles.[32] In a recent survey of the internal disclosure policies of 25 top biopharmaceutical companies, 96% reported that they had a policy committing to the sharing of summary results in academic articles or on a clinical trial registry.[33] However, in the present study, we found such commitments on the websites of only about half of the top 50 companies, suggesting that many companies are missing the opportunity to share their disclosure policies.

In contrast to the results for EFPIA/PhRMA members, but in line with those for non-member companies, a preliminary review of commitments to data transparency that was conducted for 10 non-industry sponsors of clinical trials completed in the period 2006–2015 demonstrated that only one institution referred to the disclosure of clinical trial data (Supplementary material, table S3). The requirement for EFPIA/PhRMA members to commit publicly to data disclosure is reflected in clear differences between these companies and non-members/non-industry sponsors.

Because we used data from TrialsTracker, which searches on ClinicalTrials.gov using only NCT identifiers, we made an exploratory search of alternative sources of clinical trial data for 12 clinical trials sponsored by four of the top 50 biopharmaceutical companies. Some results that were missing from ClinicalTrials.gov were found on EudraCT and company websites, suggesting that TrialsTracker was underestimating the number of trials that had published results. As recommended by the ICMJE, the inclusion of the study, NCT and/or EudraCT numbers in the abstract of publications linked to clinical trials would help to improve assessments of the disclosure of clinical trial data.[7]

Clinical trials require a high level of trust between the patient, the medical team and the trial sponsor. Central to the trust placed in the sponsor by the patient is that the results of the trial will be made publicly available. Many patients enrol in clinical trials not only in the hope of improving their own health, but also in the expectation that their participation will contribute to a better understanding of their condition and to the development of potential new treatments. Participants must weigh these potential benefits against the risk of adverse reactions, which may be serious or severe and possibly life-threatening. For their involvement in clinical trials to have meaning to the participants, trial sponsors should release all results, both positive and negative.

### Strengths and limitations

Our study was based on an evaluation of a large number of phase II–IV clinical trials from industry and non-industry sponsors over a 10-year period; however, several caveats should be considered when interpreting the results. First, it should be noted that the results obtained from TrialsTracker are subject to error in the reporting rate. As per the methods outlined in a previous article,[22] Powell-Smith and Goldacre compared their results in TrialsTracker to those from a previous manual audit of the disclosure of results from 4347 trials (performed by Chen et al.).[30] Of the 2562 trials in both analyses, 1149 were found to be reported in both, 534 were unreported by both, 497 were reported by Chen et al. but not by Powell-Smith and Goldacre, and 382 were unreported by Chen et al. but were reported by Powell-Smith and Goldacre. Thus, the total number of discordant trials (874 of 2562) represents 34.3% of the trials in both analyses. However, when analysing the results from Powell-Smith and Goldacre in comparison to those from Chen et al. more closely, 14.9% of the trials were ‘overreported’ and 19.4% were ‘underreported’, which may be interpreted as a net underestimation of the reporting rate by 4.5%; underestimation of studies performed by industry sponsors may be a particular issue because many companies only disclosed their results on their own websites. Secondly, we may not be able to generalize our findings to all sponsors and clinical trials because our analysis included sponsors of only 30 or more trials (CCW has previously calculated that sponsors of 30 studies or fewer are responsible for approximately half of all registered trials).[34] Thirdly, only two types of disclosure were included: publication in a journal and posting of results on ClinicalTrials.gov. Because publications were identified through automated searches of PubMed for NCT identifiers, identification and discoverability were limited to trials published with NCT identifiers included in the secondary source ID field of PubMed, title or abstract.[35,36] Results disclosed elsewhere (e.g. institutional websites) or published without reference to the NCT identifier could lead to the understating of disclosure rates. Fourthly, our study looked only at the disclosure of registered studies but not all studies are registered; indeed, unregistered studies seem to be less likely than registered studies to be published.[37] Finally, our analysis of publicly available disclosure policies used key word searches that focused on disclosure, so it is possible that specific publication policies were missed. Nevertheless, our findings suggest that statements related to the disclosure of results are difficult to find in many cases.

The problem of incomplete and inconsistent clinical trial disclosure remains, despite public awareness campaigns and the introduction of various policies, legislation and fines. Company-sponsored trials have been the focus of many of these activities because of their perceived commercial influence.[22] However, the present data demonstrate that results from trials sponsored by the biopharmaceutical industry are disclosed more often than those from non-industry funded studies. The results of our analysis agree with those from two recent studies that reported that industry funders disclose the results from a higher proportion of their trials than do non-industry funders.[18,27] These findings may reflect the considerable resources that commercial organizations have dedicated to clinical trial disclosure. They also suggest that the focus of future efforts to improve trial disclosure should shift towards the harmonisation of clinical trial data transparency principles to make them more easily implemented by organisations without the resources of pharma companies. This could be achieved by active discussion between, and endorsement by, all stakeholders, including clinical trial sponsors, regulatory bodies and other public bodies (e.g. WHO, ICMJE and EU Council), as well as those campaigning for increased transparency of clinical trial information. We believe that well-defined EFPIA/PhRMA joint principles could be used as a basis for the development of harmonized transparency and disclosure principles, and meanwhile should be established as an example of best practice in order to encourage consensus. Simplification of the transparency rules and regulations, the implementation of a single identifier that can be used across all registries and results databases, and improved scrutiny of compliance should extend across all aspects of clinical trials and sponsors.

Currently, transparency around clinical trial disclosure is very complex and is hindered by the absence of harmonised rules and a globally applicable platform that encompasses all registration and reporting of clinical trials for all sponsors. Such harmonisation and simplification would require a global agreement on the definition of transparency and clinical trial data disclosure, and on the process of how this should be managed in practice.

## COMPETING INTERESTS

All authors have completed the ICMJE uniform disclosure form at www.icmje.org/coi_disclosure.pdf and declare: JP, ES, JB and CCW are employees of Oxford PharmaGenesis, Oxford, UK; CCW owns shares in Oxford PharmaGenesis Holdings Ltd; AP and SB are employees of Shire International GmbH.

## FUNDING

Funding was provided by Oxford PharmaGenesis and Shire, employees of which reviewed and approved the draft text.

## ACKNOWLEDGEMENTS

The authors thank Ben Goldacre and Anna Powell-Smith for the use of the TrialsTracker tool, the patients involved in the clinical trials analysed in this article, Laura Schmidt and Simon Levy of Oxford PharmaGenesis for development of the infographic, and Christopher Rains and Valerie Philippon of Shire for reviewing the manuscript. The authors also thank Alan Thomas and Elizabeth Kinder for their review of this article from the patient perspective.

## DATA SHARING

The data presented in this article are updated from the following presentations:

Baronikova S, Purvis J, Beeso J, Southam E, Winchester C, Panayi A. Commitments to data sharing by pharmaceutical companies: the evolving environment. Presented at the 2017 European Meeting of the International Society for Medical Publication Professionals (ISMPP), London, UK, 17–18 January 2017 [poster presentation].

Baronikova S, Purvis J, Beeso J, Southam E, Winchester C, Panayi A. Commitments to data sharing by pharmaceutical companies: the evolving environment. Presented at the 2017 Annual Meeting of ISMPP, National Harbor, MD, USA, 1–3 May 2017. *Curr Med Res Opin* 2017;33(Suppl 1):7 [oral presentation].

Baronikova S, Purvis J, Beeso J, Southam E, Winchester C, Panayi A. Disclosure of results of clinical trials sponsored by pharmaceutical companies. Presented at the 2017 Meeting of the International Peer Review Congress (PRC), Chicago, IL, USA, 10–12 September 2017 [poster presentation].

Baronikova S, Purvis J, Winchester C, Southam C, Beeso J, Panayi A. Disclosure of results of clinical trials sponsored by pharmaceutical companies. Presented at the 2018 European Meeting of ISMPP, London, UK, 23–24 January 2018 [poster presentation].

This research was also presented at Evidence Live 2018, Oxford, UK, 18–19 June 2018.

This article is available as a preprint at https://www.biorxiv.org :

Slavka Baronikova, Jim Purvis, Eric Southam, Julie Beeso, Christopher Winchester, Antonia Panayi. Commitments by the biopharmaceutical industry to clinical trials transparency: the evolving environment. 2017;bioRxiv 349902; doi: https://doi.org/10.1101/349902

## Supplementary material

**Table S1.** Key elements of the ‘Final Rule’, EMA clinical data publication policy 0070 and EMA clinical trial regulation EU no. 536/2014.

|  | **FDAAA 801 the ‘Final Rule’[4]** | **EMA clinical data publication policy 0070[5]** | **EMA clinical trial regulation EU no. 536/2014[6,7]** |
| --- | --- | --- | --- |
| **Medicinal products covered** | FDA-regulated products that have not yet been approved, licensed or cleared by the FDA | Centrally authorised products only | Investigational medicinal products regardless of whether they have a marketing authorisation |
| **Clinical studies covered** | Includes all clinical trials for products as listed above | Clinical studies submitted to the agency as an MAA, Article 58, line extension or new indication, regardless of where the study was conducted | Clinical trials conducted in the EU and paediatric trials conducted outside the EU that are part of paediatric investigation plans |
| **Documents published** | Additional registration, summary results, full protocol and SAP, including more frequent updates to posted data | Clinical data overview (clinical overview, clinical summaries and CSRs) and the anonymisation report | All clinical trial-related information generated during the life cycle of a clinical trial (e.g. protocol, assessment of and decision on trial conduct, and summary of trial results including a lay summary, study reports and inspections) |
| **Publication channel** | ClinicalTrials.gov | https://clinicaldata.ema.europa.eu | Future EU portal and database (https://eudract.ema.europa.eu until that time) |
| **Effective date** | 18 January 2017 | 1 January 2015 (MAA or Article 58) or 1 July 2015 (line extension or new indication) | October 2018 |
| **Posted from** | 18 April 2018 | October 2016 | 2019 |

CSR, clinical study report; EMA, European Medicines Agency; FDA, Food and Drug Administration; FDAAA, Food and Drug Administration Amendments Act; MAA, marketing authorisation application; SAP, statistical analysis plan.

### Data analysis

A comparison of proportions (e.g. all industry, the top 50 companies and EFPIA/PhRMA members in the top 50 companies compared with non-industry clinical trial sponsors, EFPIA/PhRMA members vs non-members in the top 50 biopharmaceutical companies with statements committing to responsible data transparency) was calculated as follows (note that the comparison of clinical trial disclosure for industry and non-industry sponsors is used as an example):

- Proportion of trials disclosed by industry 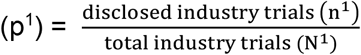
- Proportion of trials disclosed by non-industry 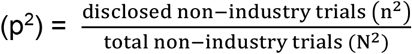
- Sample proportion 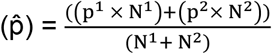
- standard error 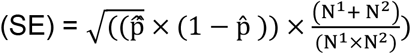
- 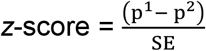

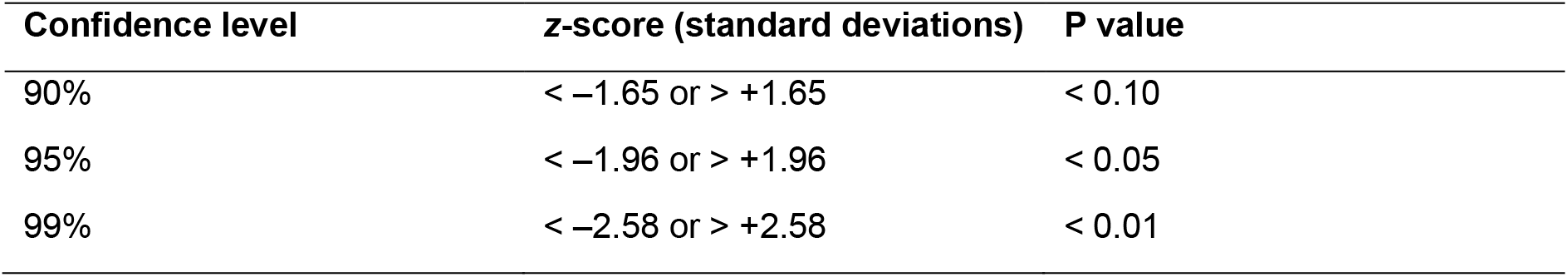
- 95% confidence interval for the difference in the proportion of disclosed clinical trials (p1 – p2):

- lower limit = (p^1^ − p^2^) − (1.96 × SE)
- upper limit = (p^1^ − p^2^) + (1.96 × SE)

The null hypothesis (H_0_) was for no difference in the proportions of disclosed trials between industry and non-industry sponsors.

**Table S2.** Comparison of disclosure rates for all industry, the top 50 biopharmaceutical companies and EFPIA/PhRMA members in the top 50 biopharmaceutical companies compared with non-industry sponsors.

| Year | Sponsor | Proportions | N | $\hat{p}$ | $1-\hat{p}$ | SE | z-value | Lower<br>95% CI | Upper<br>95% CI | Significance |
| --- | --- | --- | --- | --- | --- | --- | --- | --- | --- | --- |
| **2006** | All industry | 0.402 | 851 | 0.336 | 0.664 | 0.022 | 5.664 | 0.083 | 0.171 | Proportions significantly different at 99% |
|  | Non-industry | 0.275 | 921 |  |  |  |  |  |  |  |
| **2007** | All industry | 0.535 | 1063 | 0.401 | 0.599 | 0.020 | 11.674 | 0.192 | 0.269 | Proportions significantly different at 99% |
|  | Non-industry | 0.305 | 1478 |  |  |  |  |  |  |  |
| **2008** | All industry | 0.804 | 1361 | 0.660 | 0.340 | 0.017 | 14.892 | 0.221 | 0.288 | Proportions significantly different at 99% |
|  | Non-industry | 0.550 | 1774 |  |  |  |  |  |  |  |
| **2009** | All industry | 0.822 | 1223 | 0.659 | 0.341 | 0.017 | 15.165 | 0.226 | 0.293 | Proportions significantly different at 99% |
|  | Non-industry | 0.562 | 2063 |  |  |  |  |  |  |  |
| **2010** | All industry | 0.814 | 1108 | 0.619 | 0.381 | 0.018 | 16.356 | 0.257 | 0.327 | Proportions significantly different at 99% |
|  | Non-industry | 0.522 | 2217 |  |  |  |  |  |  |  |
| **2011** | All industry | 0.813 | 997 | 0.596 | 0.404 | 0.018 | 16.579 | 0.270 | 0.342 | Proportions significantly different at 99% |
|  | Non-industry | 0.508 | 2444 |  |  |  |  |  |  |  |
| **2012** | All industry | 0.833 | 914 | 0.589 | 0.411 | 0.019 | 17.384 | 0.291 | 0.365 | Proportions significantly different at 99% |
|  | Non-industry | 0.504 | 2630 |  |  |  |  |  |  |  |
| **2013** | All industry | 0.779 | 916 | 0.536 | 0.464 | 0.019 | 17.144 | 0.291 | 0.366 | Proportions significantly different at 99% |
|  | Non-industry | 0.451 | 2609 |  |  |  |  |  |  |  |
| **2014** | All industry | 0.809 | 763 | 0.473 | 0.527 | 0.020 | 21.033 | 0.390 | 0.470 | Proportions significantly different at 99% |
|  | Non-industry | 0.378 | 2712 |  |  |  |  |  |  |  |
| **2015** | All industry | 0.705 | 315 | 0.398 | 0.602 | 0.032 | 12.747 | 0.340 | 0.464 | Proportions significantly different at 99% |
|  | Non-industry | 0.303 | 1018 |  |  |  |  |  |  |  |
| **Overall** | All industry | 0.740 | 9511 | 0.548 | 0.452 | 0.006 | 45.626 | 0.271 | 0.295 | Proportions significantly different at 99% |
|  | Non-industry | 0.457 | 19866 |  |  |  |  |  |  |  |
|  | Top 50 | 0.760 | 6179 | 0.529 | 0.471 | 0.007 | 41.749 | 0.289 | 0.318 | Proportions significantly different at 99% |
|  | Non-industry | 0.457 | 19866 |  |  |  |  |  |  |  |
|  | EFPIA/PhRMA | 0.765 | 5785 | 0.526 | 0.474 | 0.007 | 41.267 | 0.293 | 0.322 | Proportions significantly different at 99% |
|  | Non-industry | 0.457 | 19866 |  |  |  |  |  |  |  |

CI, confidence interval; EFPIA, European Federation of Pharmaceutical Industries and Associations; $\hat{p}$ , sample proportion; PhRMA, Pharmaceutical Research and Manufacturers of America; SE, standard error.

**Table S3.** Availability of reference to transparency in clinical trial data disclosure for non-industry institutions.

| **Institution** | **Total eligible trials** | **Proportion of trials disclosed, %** | **Reference found to transparency in clinical trial data disclosure?** |
| --- | --- | --- | --- |
| **1** | 570 | 69.1 | Yes |
| **2** | 418 | 84.4 | No |
| **3** | 339 | 50.4 | No |
| **4** | 329 | 34.0 | No |
| **5** | 318 | 64.5 | No |
| **6** | 278 | 53.6 | No |
| **7** | 259 | 59.1 | No |
| **8** | 241 | 36.5 | No |
| **9** | 241 | 57.3 | No |
| **10** | 226 | 28.8 | No |

## REFERENCES

[1] The Academy of Medical Sciences. Enhancing the use of scientific evidence to judge the potential benefits and harms of medicines. 2017; Available from https://acmedsci.ac.uk/file-download/44970096 (accessed 02 Februrary 2018).

[2] World Medical Association declaration of Helsinki: ethical principles for medical research involving human subjects. JAMA, 2013;310(20):2191–4. doi: 10.1001/jama.2013.281053

[3] World Medical Association. World Medical Association declaration of Helsinki: ethical principles for medical research involving human subjects. 2018; Available from https://www.wma.net/policies-post/wma-declaration-of-helsinki-ethical-principles-for-medical-research-involving-human-subjects/ (accessed 03 May 2018).

[4] National Institutes of Health, Department of Health and Human Services. Clinical trials registration and results information submission. Final rule. 2016; 81:64981–5157. Available from https://www.ncbi.nlm.nih.gov/pubmed/27658315 (accessed 02 Februrary 2018).

[5] Regulation (EU) 536/2014 of the European Parliament and of the Council of 16 April 2014 on clinical trials on medicinal products for human use, and repealing Directive 2001/20/EC. 2014; 158:Available from https://ec.europa.eu/health/sites/health/files/files/eudralex/vol-1/reg_2014_536/reg_2014_536_en.pdf (accessed 10 April 2018).

[6] European Medicines Agency. European Medicines Agency policy on publication of clinical data for medicinal products for human use. 2015; Available from http://www.ema.europa.eu/docs/en_GB/document_library/Other/2014/10/WC500174796.pdf (accessed 09 March 2018).

[7] Panayi, A. and S. Baronikova. A new age of transparency: do we fully understand the implications? The MAP Newsletter 2017 Available from: http://ismpp-newsletter.com/2017/06/29/a-new-age-of-transparency-do-we-fully-understand-the-implications/ (accessed 23 March 2018).

[8] Slovakian State Institute for Drug Control. Available from: https://www.sukl.sk/en/clinical-trials/scope-of-activities?page_id=4021 (accessed 30 April 2018).

[9] Netherlands Trial Registry. Available from: http://www.trialregister.nl/trialreg/index.asp (accessed 30 April 2018).

[10] Japan Primary Registries Network. Available from: http://rctportal.niph.go.jp/en/ (accessed 30 April 2018).

[11] L’Agence nationale de sécurité du médicament et des produits de santé. Available from: http://ansm.sante.fr/ (accessed 30 April 2018).

[12] World Health Organization. International standards for clinical trial registries. 2012; Available from http://apps.who.int/iris/bitstream/10665/76705/1/9789241504294_eng.pdf (accessed 09 March 2018).

[13] International Committee of Medical Journal Editors (ICMJE). Recommendations for the conduct, reporting, editing, and publication of scholarly work in medical journals. 2016; Available from http://icmje.acponline.org/news-and-editorials/icmje-recommendations_annotated_dec14.pdf (accessed 09 March 2018).

[14] Battisti, W.P., E. Wager, L. Baltzer, et al., Good publication practice for communicating company-sponsored medical research: GPP3. Ann Intern Med, 2015;163(6):461–4. doi: 10.7326/M15-0288

[15] GlaxoSmithKline. Clinical study register. Available from: https://www.gsk-clinicalstudyregister.com/ (accessed 21 May 2018).

[16] Bayer AG. Clinical study finder. Available from http://pharma.bayer.com/en/innovation-partnering/clinical-trials/trial-finder/ (accessed 21 May 2018).

[17] Ehrhardt, S., L.J. Appel, and C.L. Meinert, Trends in National Institutes of Health funding for clinical trials registered in ClinicalTrials.gov. JAMA, 2015;314(23):2566–2567. doi:

[18] Zwierzyna, M., M. Davies, A.D. Hingorani, et al., Clinical trial design and dissemination: comprehensive analysis of clinicaltrials.gov and PubMed data since 2005. BMJ, 2018;361. doi: 10.1136/bmj.k2130

[19] European Federation of Pharmaceutical Industries and Associations and Pharmaceutical Research and Manufacturers of America. EFPIA and PhRMA joint principles for responsible clinical trial data sharing. 2013; Available from http://phrma-docs.phrma.org/sites/default/files/pdf/PhRMAPrinciplesForResponsibleClinicalTrialDataSharing.pdf (accessed 12 February 2018).

[20] Looney, W. Pharm exec’s top 50 companies 2016. 24 November 2017. Available from: http://www.pharmexec.com/2016-pharm-exec-50 (accessed 13 December 2017).

[21] Zeldman, J., Taking Your Talent to the Web: A Guide for the Transitioning Designer, ed. S. Champeon. 2001, Indianapolis, USA: New Riders.

[22] Powell-Smith, A. and B. Goldacre, The TrialsTracker: automated ongoing monitoring of failure to share clinical trial results by all major companies and research institutions. F1000Res, 2016;5:2629. doi: 10.12688/f1000research.10010.1

[23] Porter, T. and R. Miller, Investigating the three-click rule: a pilot study. MWAIS Proceedings. 2., 2016. doi:

[24] Mooney, L.A. and L. Fay, Cross-sectional study of Pfizer-sponsored clinical trials: assessment of time to publication and publication history. BMJ Open, 2016;6(7):e012362. doi: 10.1136/bmjopen-2016-012362

[25] Evoniuk, G., B. Mansi, B. DeCastro, et al., Impact of study outcome on submission and acceptance metrics for peer reviewed medical journals: six year retrospective review of all completed GlaxoSmithKline human drug research studies. BMJ, 2017;357:j1726. doi: 10.1136/bmj.j1726

[26] Ross, J.S., M. Mocanu, J.F. Lampropulos, et al., Time to publication among completed clinical trials. JAMA Intern Med, 2013;173(9):825–8. doi: 10.1001/jamainternmed.2013.136

[27] Goldacre, B., N.J. DeVito, C. Heneghan, et al., Compliance with requirement to report results on the EU Clinical Trials Register: cohort study and web resource. BMJ, 2018;362:k3218. doi: 10.1136/bmj.k3218

[28] Miller, J.E., M. Wilenzick, N. Ritcey, et al., Measuring clinical trial transparency: an empirical analysis of newly approved drugs and large pharmaceutical companies. BMJ Open, 2017;7(12). doi: 10.1136/bmjopen-2017-017917

[29] Deane, B.R. and S. Porkess, Clinical trial transparency update: an assessment of the disclosure of results of company-sponsored trials associated with new medicines approved in Europe in 2014. Curr Med Res Opin, 2018;1–5. doi: 10.1080/03007995.2017.1415057

[30] Chen, R., N.R. Desai, J.S. Ross, et al., Publication and reporting of clinical trial results: cross sectional analysis across academic medical centers. BMJ, 2016;352:i637. doi: 10.1136/bmj.i637

[31] Tompson, A.C., S. Petit-Zeman, B. Goldacre, et al., Getting our house in order: an audit of the registration and publication of clinical trials supported by the National Institute for Health Research Oxford Biomedical Research Centre and the Musculoskeletal Biomedical Research Unit. BMJ Open, 2016;6(3):e009285. doi: 10.1136/bmjopen-2015- 009285

[32] EFPIA and PhRMA. EFPIA-PhRMA principles for responsible clinical trial data sharing: report on the 2016 member company survey. 2016; Available from https://www.efpia.eu/media/288603/efpia-phrma-report-on-the-2016-member-company-survey-on-the-joint-principles-for-responsible-clinical-trial-data-sharing.pdf (accessed 28 March 2018).

[33] Goldacre, B., S. Lane, K.R. Mahtani, et al., Pharmaceutical companies’ policies on access to trial data, results, and methods: audit study. BMJ, 2017;358:j3334. doi: 10.1136/bmj.j3334

[34] Winchester, C., Reader comment on: The TrialsTracker: automated ongoing monitoring of failure to share clinical trial results by all major companies and research institutions [version 1; referees: 2 approved]. F1000Research, 2016;5:2629. doi:

[35] Coens, C., J. Bogaerts, and L. Collette, Comment on the “TrialsTracker: automated ongoing monitoring of failure to share clinical trial results by all major companies and research institutions” [version 1; referees: 1 approved, 1 approved with reservations]. F1000Research, 2017;6:71. doi: doi: 10.12688/f1000research.10503.1

[36] Mooney, L.A., J.F. Michalski, and L. Fay, Presence of a unique trial identifier in the abstracts of industry-sponsored manuscripts, in Peer Review Congress (PRC). Chicago, IL, USA.

[37] Chan, A.W., A. Pello, J. Kitchen, et al., Association of trial registration with reporting of primary outcomes in protocols and publications. JAMA, 2017;318(17):1709–1711. doi: 10.1001/jama.2017.13001

